# Matching Corticospinal and Cortical Excitability along the motor cortex: TMS-EMG-EEG study

**DOI:** 10.64898/2026.11.03.763103

**Authors:** Maria Nazarova, Macey Higdon, Joel Rouste, Tuomas Mutanen, Vadim Nikulin, Elena Ukharova, Selja Vaalto, Risto J. Ilmoniemi, Pantelis Lioumis, Hanna Renvall

## Abstract

There is still no consensus on how to choose intensities for transcranial magnetic stimulation (TMS). The resting motor threshold (RMT)—the intensity needed to evoke muscle contractions in the contralateral limb muscles—is a robust muscle-specific index of corticospinal excitability. RMT from one of the hand muscles is commonly used as a reference for TMS neuromodulation interventions and TMS-electroencephalography (TMS-EEG) studies. However, it only provides an indirect measure of cortical excitability, and even for the primary motor cortex (M1), we do not understand how cortical and corticospinal excitability correspond to each other. Indeed, it remains unclear whether cortical responses to TMS follow similar location-dependent patterns along the M1 as RMTs, and whether subthreshold stimulation can reliably engage cortical circuits relevant to motor output. Combining TMS with electromyography (EMG) and EEG—TMS-EMG-EEG—enables simultaneous assessment of corticospinal and cortical excitability to answer these questions.

We investigated how cortical excitability thresholds (CETs, measured with EEG), relate to corticospinal excitability thresholds (RMTs, measured with EMG) at the hand, arm, and leg hotspots along the M1 in 24 healthy subjects. RMTs and CETs were strongly correlated, and both increased from hand to arm to leg. CETs were generally lower than RMTs, indicating that subthreshold stimulation was enough to evoke robust early TMS evoked potentials (TEPs). Within subjects, CETs differed from the corresponding RMTs at all three M1 hotspots, and this difference increased from hand to leg representations.

The findings of the present study argue against using a default hand-muscle RMT to set cortical excitability dosage along the M1 and beyond. Online TMS–EEG can enable real-time dosing and could improve stimulation precision, efficiency, and generalizability, but the site-specific variability calls for adaptive control.

**Dictionary:** *TMS-EMG-EEG:* A multimodal neurophysiological approach, combining EMG and EEG recording during TMS, to assess corticospinal and cortical excitability simultaneously.

*Corticospinal Excitability:* Responsiveness of the corticospinal motor pathway to stimulation, reflecting the combined excitability of cortical and spinal motor circuits.

*Cortical Excitability:* Responsiveness of cortical neuronal circuits to external stimulation, including electrical and magnetic stimulation.

*EMG-EEG Compound Hotspot:* The stimulation site that elicited clear and consistent early TEPs and reliable MEPs for the target muscle.

*MEP (motor evoked potential):* Electrical activity recorded from the peripheral muscle via EMG, in response to TMS stimuli.

*RMT (resting motor threshold):* A measure of corticospinal excitability.

*TEP (TMS-evoked potential):* Electrical potentials recorded from the scalp via EEG, in response to TMS stimuli.

*CET (cortical excitability threshold):* The early peak-to-peak deflection at an electrode near the stimulation site, quantified in real-time.

## Introduction

It is well established that corticospinal excitability, reflected by transcranial magnetic stimulation (TMS)-derived resting motor thresholds (RMTs), varies systematically across primary motor cortex (M1), with RMTs increasing from hand to arm to leg hotspots (Chen, Tam, *et al*., 1998; Rossini *et al*., 2015; Nazarova, Novikov, *et al*., 2021; Yuasa *et al*., 2022; Koponen *et al*., 2024). This gradient is thought to be connected to sodium channel antagonists at the cortical level (Ziemann *et al*., 1996; Rocchi *et al*., 2025). At the same time, differences in cortico-spinal connectivity likely also contribute (Brouwer and Ashby, 1990; Palmer and Ashby, 1992; Groppa *et al*., 2012). Studies using simultaneous TMS and electroencephalography (TMS–EEG) have shown that TMS cortical excitability differs across neighboring areas with spatial resolution on the order of ∼1 cm (Ruohonen and Ilmoniemi, 1998; Passera *et al*., 2022). Most TMS–EEG studies use stimulation intensities relative to hand-muscle RMT, with >70% using suprathreshold values (Beck *et al*., 2024). The dosing choice is challenging: TEP amplitudes may be only 2-3 μV, i.e., making their detection highly dependent on signal-to-noise ratio (Hernandez-Pavon *et al*., 2023; Parmigiani *et al*., 2023). At the same time, high TMS intensities can increase sensory/afferent contamination and discomfort (Nikouline, Ruohonen and Ilmoniemi, 1999; Conde *et al*., 2019). The real-time TMS–EEG approach (Iscan *et al*., 2016; Casarotto *et al*., 2022; Tervo *et al*., 2022; Ukharova *et al*., 2025, 2026; *eLife*, 2025) provides real-time estimates of local cortical excitability by visualizing TEPs during stimulation and quantifying the early peak-to-peak deflection at an electrode near the stimulation site (Casarotto *et al*., 2022; D’Ambrosio *et al*., 2022; Rissanen *et al*., 2026). However, how this cortical excitability threshold (CET) relates to RMTs across M1 remains unknown. It is also unclear whether EEG-based dosing is stable over the several hours typical for TMS-EEG measurements.

Our aims were twofold: 1) to test whether the known corticospinal RMT gradient is cortically mirrored by the early TEP deflection (CET), and 2) to assess the within-session stability of such CET. This work links local cortical signals (EEG), descending corticospinal output (electromyography; EMG), and, prospectively, network-level connectivity measures for optimal TMS dosing in an integrated framework. In the long term, dose–response maps integrating cortical and corticospinal signatures along the M1 could support diagnostic and therapeutic applications in patients with motor impairments associated with altered brain excitability (Turton *et al*., 1996; Bütefisch *et al*., 2003; Beck and Hallett, 2011; Nazarova, Kulikova, *et al*., 2021; Harquel *et al*., 2024; Ziemann *et al*., 2026), complementing recent efforts toward individualized TMS targeting (Lomi *et al*., 2026; Lioumis *et al*., 2025).

## 2. Methods

### 2.1 Subjects

24 healthy right-handed individuals (11 females; age range 21-46 years, mean 30 +-6 years) participated in the study. All subjects provided written informed consent in accordance with the Declaration of Helsinki after a favorable statement from the HUS Regional Committee on Medical Research Ethics (HUS/1198/2016). Exclusion criteria included contraindications for TMS or MRI, any known neurological or psychiatric disorders, and taking medication affecting the central nervous system. No TMS-related adverse events were reported during the study.

### 2.2 Experimental Setup

All the experiments were performed at Aalto TMS Laboratory at the Department of Neuroscience and Biomedical Engineering, Aalto University School of Science.

#### 2.2.1 Transcranial Magnetic Stimulation (TMS)

TMS was performed using MRI-navigated TMS (Nexstim NBT 2.2.4, Nexstim Plc, Finland) with a 70 mm figure-of-eight coil. We used individual structural T1-weighted anatomical MRIs for navigation (1 mm slice thickness, MAGNETOME Skyra 3 T, Siemens, Germany). Peeling depth in the neuronavigation system was set between 20-25 mm for optimal visualization of the M1. Biphasic pulses with an interstimulus interval of 2000 ms and uniformly distributed random jitter of 2-2.4 ms, with a recharge delay of 900 ms, were used to avoid potential recharge artifacts in EEG recordings (Ukharova *et al*., 2025). During stimulation, subjects sat comfortably in an armchair with arm and head rests, while their upper limbs rested in a pronated position on a C-shaped pillow (Figure 1). The subjects were instructed to gaze at a fixation cross positioned in front of them at eye level. Stimulation intensity values were recorded in both % of maximum stimulator output (% MSO) and E-field strength (V/m) according to Nexstim’s 3-D MRI head model (Hannula and Ilmoniemi, 2017).

**Figure 1:**
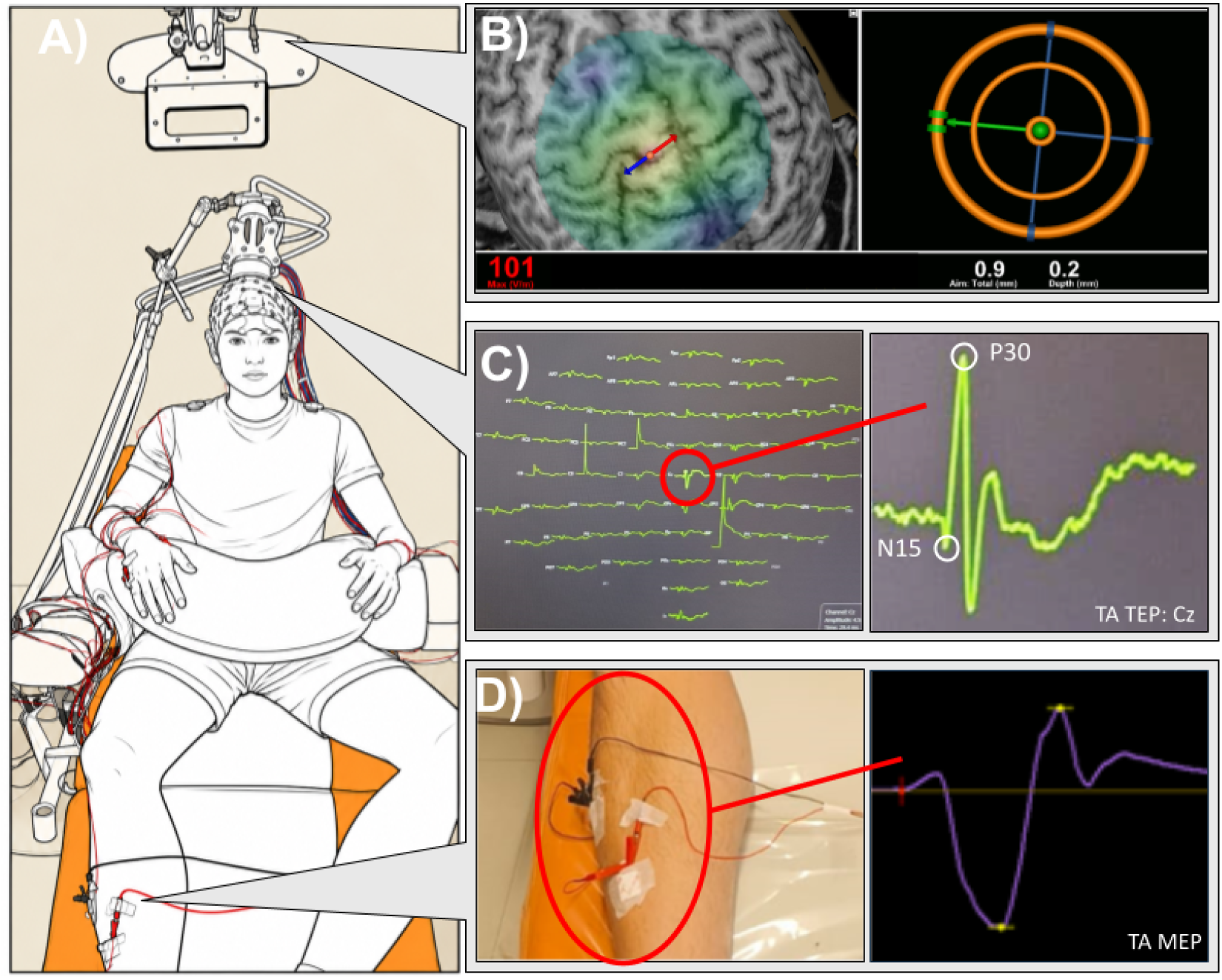
TMS-EMG-EEG Experimental Setup. (A) EEG cap fixed to the head, with the TMS coil positioned over the TA hotspot. One can see bipolar surface EMG electrodes on tibialis anterior (TA), biceps brachii (BB), abductor pollicis brevis (APB), and abductor digiti minimi (ADM). (B) Nexstim infrared camera for navigation and an example 3D MRI reconstruction in the Nexstim software, with the stimulation target shown at the TA hotspot. (C) Example array of ∼20 TEPs averaged, displayed on the real-time TEP (rt-TEP) monitor during stimulation of the TA hotspot, with a zoomed-in view of a corresponding TA TEP from electrode Cz. (D) Exemplary EMG electrode placement and the corresponding TA motor-evoked potential (MEP). Image edited using ChatGPT (OpenAI, 2026) to create an outline of the setup.

#### 2.2.2 Electromyography (EMG) recording

MEPs were obtained from 1) two intrinsic hand muscles [abductor pollicis brevis (APB) and abductor digiti minimi (ADM)], 2) one arm muscle [biceps brachii (BB)], and 3) one leg muscle [tibialis anterior (TA)] on the right side (Figure 1a). MEPs from the ADM muscle were monitored to trace the cortical somatotopy within the hand (Fig. 1a). Bipolar surface electrodes (Red Dot, 3M, USA) were placed in a belly-tendon montage using touch-proof connectors (Cephalon, Denmark), connected to both the Nexstim integrated EMG for online MEP analysis and the Bittium NeurOne system (Bittium Biosignals, Finland) for offline analysis alongside EEG (Figure 1d). EMG was continuously monitored visually, and if it exceeded 20 μV peak-to-peak, subjects were asked to relax or slightly change their arm and hand position.

#### 2.2.3 Electroencephalography (EEG) recording

EEG was recorded using a TMS-compatible EEG amplifier (NeurOne, Bittium Biosignals, Finland) with a 64-electrode EEG cap with multitrode passive electrodes (BrainCap for TMS, Easycap, Germany). During the acquisition, EEG signals were low-pass filtered at 1250 Hz with a sampling frequency of 5 kHz. The reference and ground electrodes were placed on the right mastoid and cheek, respectively. During the recordings, the electrode impedances remained below 5 kΩ, with typical values around 2 kΩ. We monitored online TEPs through a real-time graphical user interface, based on (Casarotto *et al*., 2022). We utilized TAAC toolbox (Russo *et al*., 2022) to generate temporally and spectrally matched, TMS-synchronized masking noise, delivered through pneumatic earplugs, to reduce TMS-evoked auditory responses. We individualized the sound level by observing online TEPs after approximately 20 test stimuli, with the coil positioned ∼8 cm from the head and using near-maximal stimulator output (MSO) stimuli (Ukharova *et al*., 2025). Noise masking was considered adequate when no discernible N100 component (Nikouline, Ruohonen and Ilmoniemi, 1999) was observed in the averaged TEP under any electrode and when subjects self-reported that they could not hear the coil click.

### 2.3 Experimental Procedure

#### 2.3.1 M1 TMS-EMG: rough motor mapping

TMS-EMG rough motor mapping was performed prior to searching the MEP/TEP threshold, similarly to our earlier approach (Nazarova, Novikov, *et al*., 2021). The EEG cap was fitted (but not yet prepped) to maintain a consistent coil-to-scalp distance. We identified four to five tentative “MEP hotspots” along M1 (defined as the coil positions that produced the highest MEP amplitudes in the target muscle) for APB (hand), BB (arm), and TA (leg) muscles.

#### 2.3.2 M1 TMS-EMG-EEG: compound hotspot mapping

Following TMS-EMG rough motor mapping, the EEG cap was prepped to perform simultaneous TMS-EMG-EEG and to identify a compound MEP-TEP hotspot for each of the three M1 sites (hand, arm, leg). We defined the **compound EMG-EEG hotspot** as the stimulation site that elicited clear and consistent early TEPs and reliable MEPs for the target muscle. This compound hotspot was determined for each site by the following: Approximately 20 stimuli were given at the first tentative MEP hotspot while TEPs were observed online. The hotspot was rejected if stimulation elicited a significant artifact, typically caused by scalp muscle activation (Figure 2b). If there were no artifacts, at least 2 other tentative MEP hotspots were tested to identify the hotspot with the largest early TEP peak-to-peak amplitude (at 8–35 ms latency). After the compound hotspot was selected, we determined the TEP threshold, i.e. “cortical excitability threshold” (CET), defined as the stimulation level that evokes an 8 µV deflection at the earliest visible TEP deflection of at least one electrode near the stimulation site these electrodes varied considerably: for APB hotspot there were 9, for BB hotspot −11, and for TA hotspot - 8 possible electrodes (see Figure S1 in Supplementary Information). To find the CET, 20 stimuli were given in 1-2% MSO increments until the early TEP deflection reached ≥ 8 µV average amplitude. We have chosen 8 µV as an operational compromise value, considering the use of the values from 6 to 10 µV in the earlier works (Casarotto *et al*., 2022; D’Ambrosio *et al*., 2022). After determining the CET, we measured the RMT using an adaptive threshold hunting algorithm (Julkunen, 2019). These steps were repeated for each compound hotspot. The distance between each compound hotspot was at least 1 cm (APB–BB 10–21 mm, BB–TA 17–30 mm, APB–TA 27–50 mm) (see Figure S2 in Supplementary Information). In 4 subjects, stimulating the BB hotspot caused a large muscle artifact, so we did not collect data on BB stimulation in those subjects.

**Figure 2:**
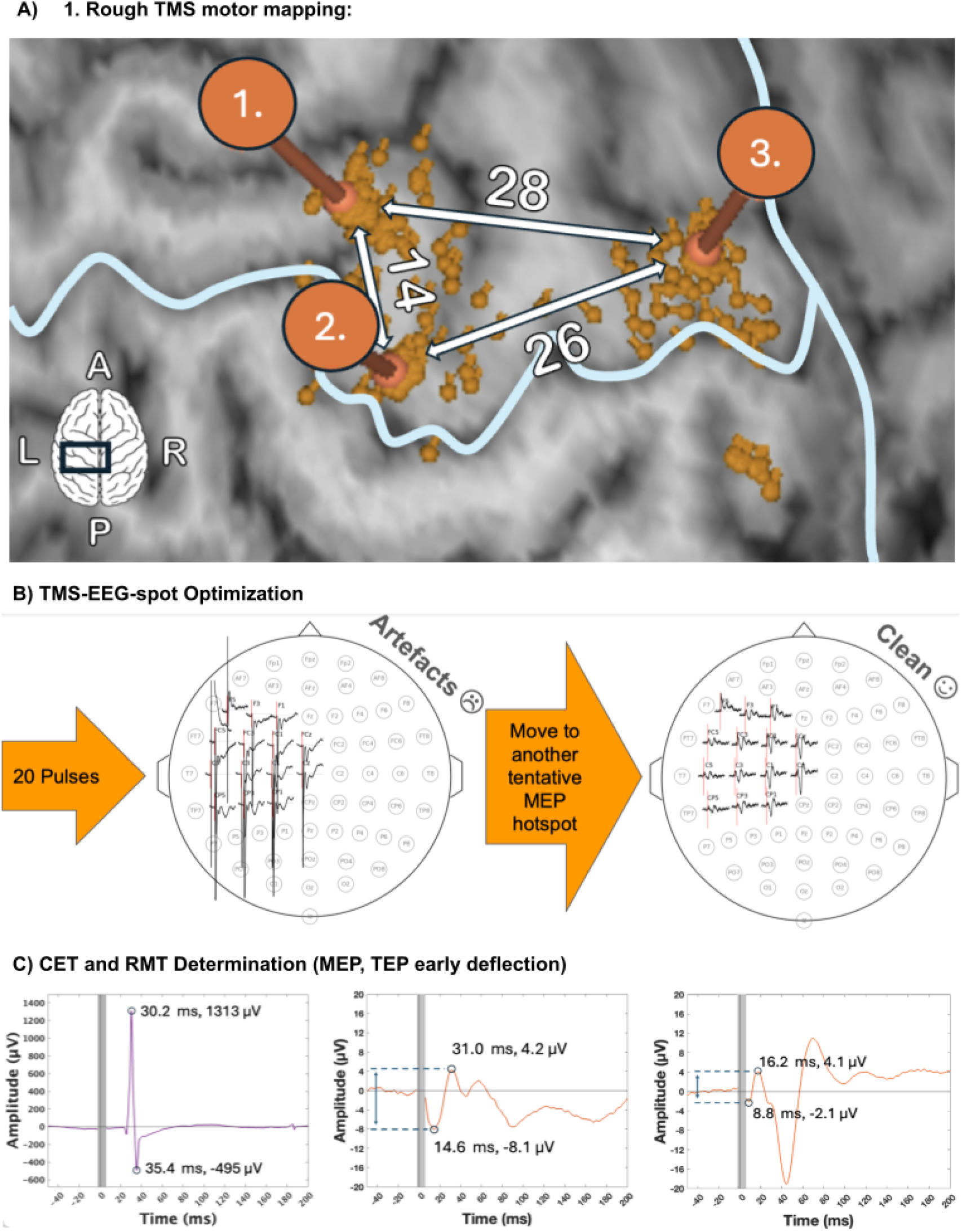
TMS-EMG-EEG Experimental Procedure. A) Exemplary rough motor mapping and corresponding compound hotspots (shown by the three orange circles) (1) APB, (2) BB, and (3) TA, with Euclidean distances between them in mm. B) Compound hotspot search procedure. Example of artifacts in the electrode of interest after averaging ∼20 TEPs, which led to testing an alternative tentative MEP hotspot. C) Left: Example of MEP (APB hotspot, APB muscle). Middle: Example of a TEP (BB hotspot) where the measured peak-to-peak amplitudes closely correspond to the N15 and P30. Right: Example of a TEP (TA hotspot) where the measured peak-to-peak amplitudes was earlier than the N15-P30.

The analyzed experiment consisted of three measurement sessions (8 additional non-repetitive TMS-EEG sessions were performed during the experiment, but they are out of the scope of the current article). In each session, 120 stimuli were delivered at CET intensity per compound hotspot (APB, TA, and BB) at rest. Subjects were reminded to stay alert and fixate on the cross; researchers monitored that eyes remained open throughout the sessions. For simplicity, the term hotspot from this point on will refer to the compound hotspot.

### 2.4 Data Analysis

#### 2.4.1 EEG Analysis

The EEG data were minimally preprocessed using a custom MATLAB script (R2024b, v9.8, The MathWorks, Inc., Natick, USA) based on EEGLAB (Delorme and Makeig, 2004), and the TMS-EEG signal analyzer (TESA) toolbox (Rogasch *et al*., 2017). Continuous EEG data were epoched from −1500 ms to 1500 ms and baseline corrected from −150 ms to −10 ms relative to the TMS trigger. TMS pulse artifact (from −2 ms to 6-10 ms) was removed based on visual inspection, depending on the length of the muscle or decay artifact. The cut segment length was kept the same for each condition within the same hotspot. Bad electrodes (e.g., big decay artifact) were removed based on visual inspection of channel spectra and raw EEG. Bad epochs (significant ocular or movement artifacts) were removed based on raw data time series. This resulted in an average 112 ± 8 epochs being included in the analysis. We used FastICA (Bingham and Hyvärinen, 2000) only to remove remaining blink artifacts. After blink removal, we re-referenced the data to a common reference and applied a low-pass filter (zero-phase, non-causal) with a cut-off frequency of 200 Hz. In 10 measurements across 6 subjects, we had to remove the electrode that was used for CET determination due to remaining artifacts (Figure 2). We visually inspected peak-to-peak amplitudes to identify the earliest positive and negative TEP deflections (usually N15 and P30, though sometimes earlier TEP deflections were also observed; see Figure 2): We used these early TEP deflections at each hotspot as the measure of cortical excitability. To assess the stability of the cortical excitability throughout the measurement sessions, we measured the early TEP deflection in bins of 20 consecutive epochs. Statistical analyses were performed on the first five bins, corresponding to epochs 1–100 of the preprocessed data. The final bin (epochs 101–120) was excluded from the analysis because the number of remaining epochs varied across measurement sessions due to epoch rejection during preprocessing. We assessed the stability of the early TEP deflections by comparing their amplitudes to those measured in the subsequent TMS-EEG session 30–90 minutes later. We also examined the bins against each other to investigate short-scale dynamics of the early TEPs.

#### 2.4.2 EMG Analysis

EMG signal was recorded with the Bittium NeurOne system and processed in EEGLAB with a 50-Hz notch and 10-1000 Hz bandpass filter. MEPs were detected automatically within predefined latency windows (18-25 ms for APB and ADM, 10-20 ms for BB, and 25-35 ms for TA) and required a peak-to-peak amplitude difference > 50 μV. Epochs showing > 50 μV peak-to-peak preactivation 500 ms before the stimulus in the targeted muscle were omitted. In cases where full relaxation was not achieved (n=7, one session per subject), we accounted for preactivation as follows: If preactivation occurred in a non-targeted muscle, the MEP detection threshold for that electrode was increased from 50μV to 200μV. This adjustment was done to minimize the exclusion of epochs from the dataset. All automatically detected results were subsequently confirmed by visual inspection. The MEP rate (%) was defined as the proportion of epochs containing MEPs that occurred within the latency windows described above and met the amplitude threshold criteria. The MEP rate was computed for all four muscles in all measurements. Sessions with less than 50 epochs after preprocessing were not analyzed (n=3).

### 2.5 Statistical Analyses

We conducted statistical comparisons using custom-made Python scripts and GraphPad Prism 11 software. We tested all results for normality using the Shapiro-Wilk test. When normality was rejected (RMT and CET thresholds, p < 0.05), we used non-parametric statistical tests. We compared the dependent variables of interest using the Friedman test for multiple comparisons, with Kendall’s W ranging from 0 (no concordance) to 1 (perfect concordance). If overall differences (uncorrected p < 0.05) were detected, we performed post-hoc Wilcoxon signed-rank tests, adjusting p-values for multiple comparisons using the Benjamini-Hochberg false discovery rate (FDR)(Benjamini and Hochberg, 1995). Effect size was tracked by calculating the Hedges’ g measure of effect size. To assess a possible shared source of variability between corticospinal and cortical measures, we performed principal component analysis (PCA) on RMTs and CETs across hotspots.

## 3. Results

### 3.1. Both corticospinal (RMT) and cortical (CET) thresholds increase from hand to arm to leg representations, with a shallower gradient for cortical thresholds

We first examined whether the well-known medio-lateral gradient in corticospinal excitability is mirrored by a corresponding gradient in cortical excitability. Both RMT and CET increased systematically from the hand to the arm and leg representations (RMT: W=0.68; CET: W=0.60; Figure 3). This pattern was observed both when thresholds were expressed as estimated EF values or as % MSO (all results are expressed in V/m; %MSO results can be found in Supplementary Information) The medio-lateral gradient was less pronounced for the cortical thresholds than for the corticospinal RMTs (median RMT: APB = 61 V/m, BB = 76 V/m, TA = 95.5 V/m; median CET: APB = 54.5 V/m, BB = 61.5 V/m, TA = 73 V/m; Figure 3; Descriptive statistics in Supplementary Information). While hand thresholds were always lower than leg thresholds, the gradient was not absolute. We observed four exceptions for RMTs (2 cases with BB RMT > APB RMT, 2 cases with BB RMT > TA RMT) and seven for CETs (4 cases with APB CET >= BB CET, 3 cases with BB CET > TA CET). In no case were leg thresholds lower than or equal to hand thresholds.

**Figure 3:**
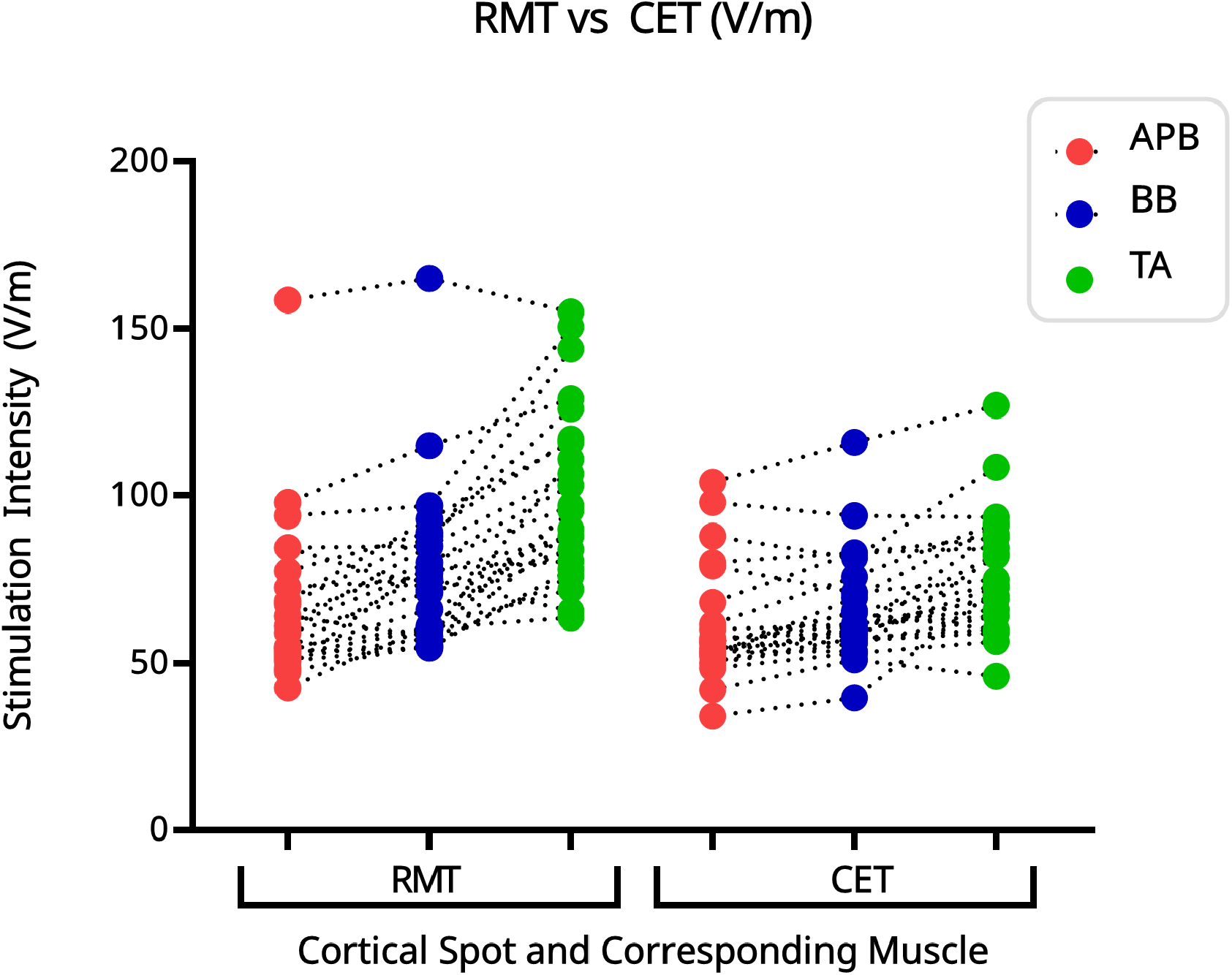
RMTs vs. CETs (V/m) gradient at the hand (APB; red), arm (BB; blue), and leg (TA; green) hotspots on M1.

All six RMT and CET thresholds were positively correlated (Table 1). We found high correlations for homotopic pairs, among thresholds within upper-limb representations, and between the biceps and TA hotspots when thresholds were obtained using the same thresholding method. The remaining cross-limb associations showed moderate correlations.

**Table 1:** Median and Correlation Matrix for RMT and CET. hand-arm-leg hotspots across the M1. Spearman’s R and P Values (***p<0.001, ***p<0.0001, FDR corrected).

| Variable | *Median* | APB RMT | BB RMT | TA RMT | APB CET | BB CET | TA CET |
|---|---|---|---|---|---|---|---|
| APB RMT | 61 | — |  |  |  |  |  |
| BB RMT | 76 | .76*** | — |  |  |  |  |
| TA RMT | 95.5 | .59** | .72*** | — |  |  |  |
| APB CET | 54.5 | .87*** | .69*** | .57** | — |  |  |
| BB CET | 61.5 | .77*** | .68*** | .65** | .85*** | — |  |
| TA CET | 72.75 | .57** | .57** | .72*** | 0.62** | .71*** | — |

In PCA analysis, the first principal component explained 85% of the variance for RMTs and 88% for CETs. First principal components were highly correlated (Spearman’s r = 0.80, p < 0.001).

### 3.2. Hotspot specific subthreshold intensities elicit clear TEPs in M1, and CET and RMT is spot-specific

Across all hotspots, CETs were generally lower than RMTs (Figure 3 and 4).

**Figure 4:**
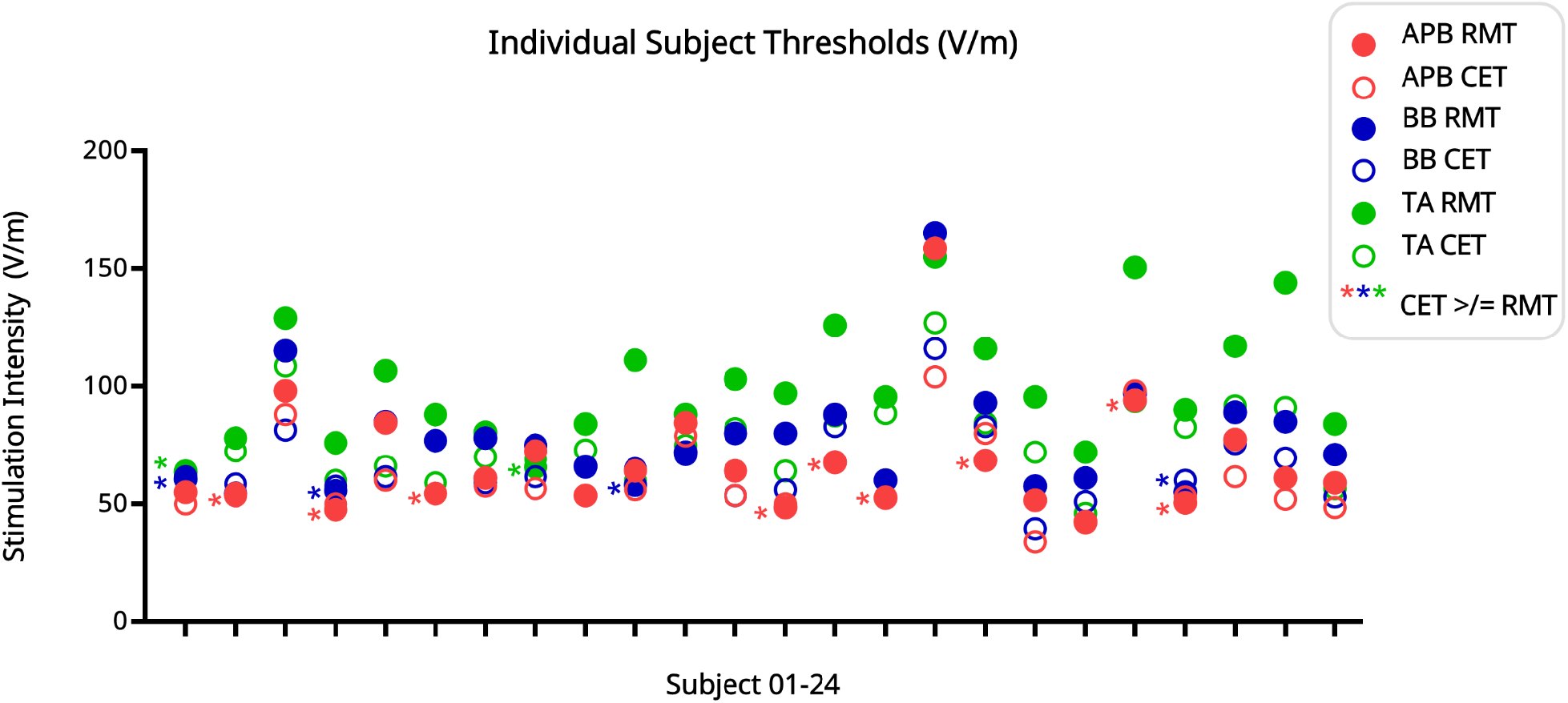
**Subject Specific RMTs** (filled circles) **vs. CETs (V/m)** (empty circles) at hand (APB, red), arm (BB, blue) and leg (TA, green) hotspots on the M1. Stars (*) on the left side of the circles indicate those cases where the CET was greater than or equal to RMT at the same hotspot (according to the hotspot color coding).

On average, the median difference between CET and RMT (RMT-CET=threshold difference in V/m) decreased latero-medially: 7.26 V/m in the APB hotspot, 11.49 V/m in the BB hotspot, and 23.59 V/m in TA hotspot (see Table S3 in Supplementary Information). Nonetheless, in 16 cases, CETs exceeded RMTs (9 cases for APB, 5 cases for BB, and 2 cases for TA (Figures 3-4). In 3 subjects, this occurred at both upper-limb hotspots.

RMT is a muscle-specific measure, defined as the lowest intensity that evokes around 50% of ≥50 μV MEPs in this resting muscle (Rossi *et al*., 2021; Nazarova and Asmolova, 2022). Consequently, intermittent MEPs can occur at subthreshold intensities below RMT. Figure 5 represents the association between the homotopic MEP rate (APB muscle for APB hotspot, BB muscle for BB hotspot, TA muscle for TA hotspot) and early TEP-average amplitude: most TEPs could be observed when stimulation was below RMT but still evoked intermittent MEPs.

**Figure 5:**
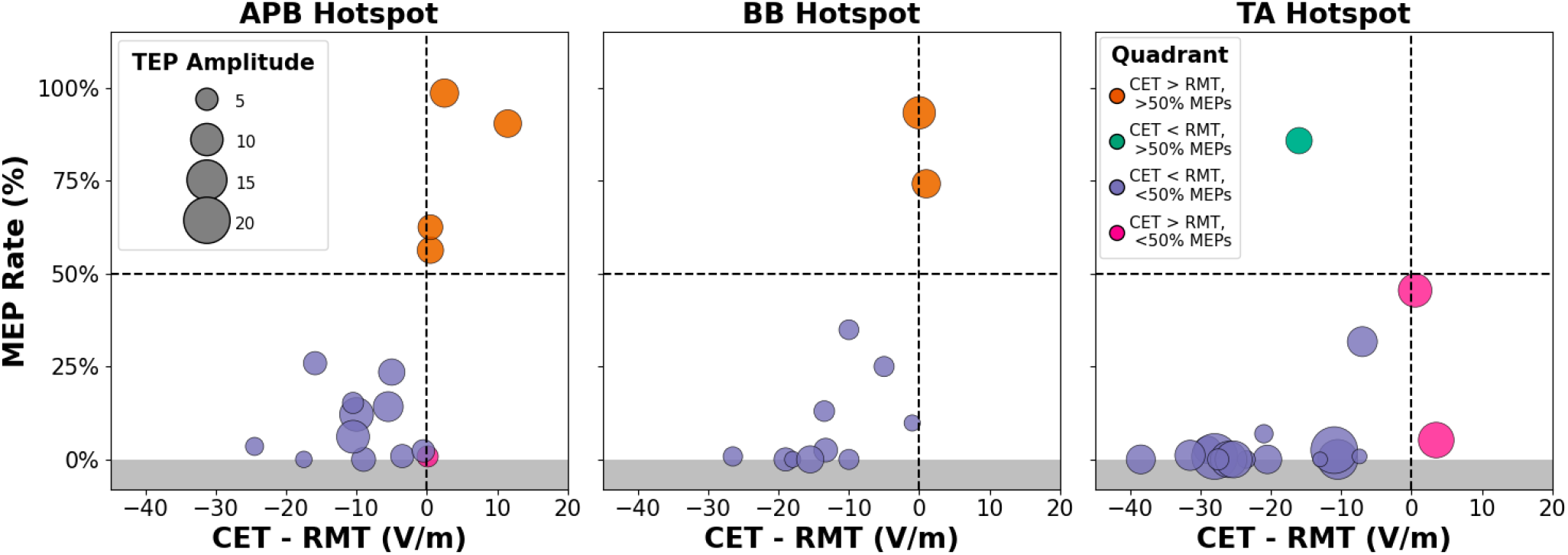
MEP rate at each hotspot. MEP rate presented as % of MEPs ≥ 50μV across 120 epochs, plotted against the stimulation intensity difference between CET and RMT (CET−RMT). Circle diameters represent the TEP amplitudes. The horizontal dashed line (x-axis) represents the 50% MEP rate and the vertical dashed line (y-axis) the intensity where CET and RMT intensities are equal.

### 3.3. Subthreshold TMS-EMG excitability is muscle-specific

We further assessed the muscle specificity of subthreshold corticospinal excitability at all three hotspots along the M1 (Figure 6). As expected, stimulation at the APB hotspot produced the highest MEP rates in APB muscle. In contrast, at the BB hotspot, intrinsic hand muscles showed MEP rates comparable to BB muscle.

**Figure 6:**
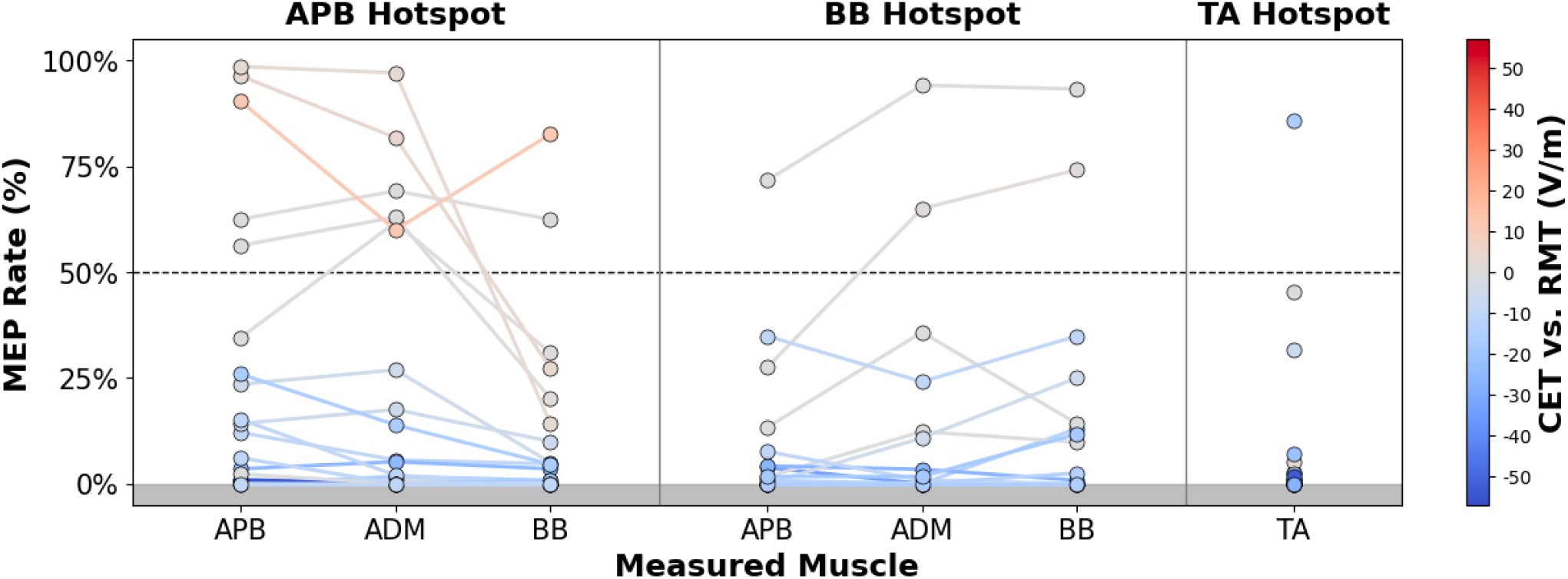
Distribution of MEP rates (%) across muscles for each hotspot. Colour indicates the stimulation intensity difference between CET and RMT (CET−RMT). For the TA hotspot, only the TA MEPs are shown, as TA hotspot stimulation did not evoke MEPs in the upper limb muscles, and the TA EMG electrode is omitted from the APB and BB cortical hotspot stimulation results for the same reason.

### 3.4. Limitations in CET Determination

The stability of early TEP deflection amplitude was hotspot-specific: at the APB hotspot, the amplitudes decreased statistically significantly compared to the target 8 µV (mean 8 µV vs. 6.1 µV, V=15.0, p = 0.001; One-sample Wilcoxon signed-rank test), at the BB hotspot the decrease was not statistically significant. At the TA hotspot, values varied widely among subjects (mean = 8.9 µV) (Figure 7).

**Figure 7:**
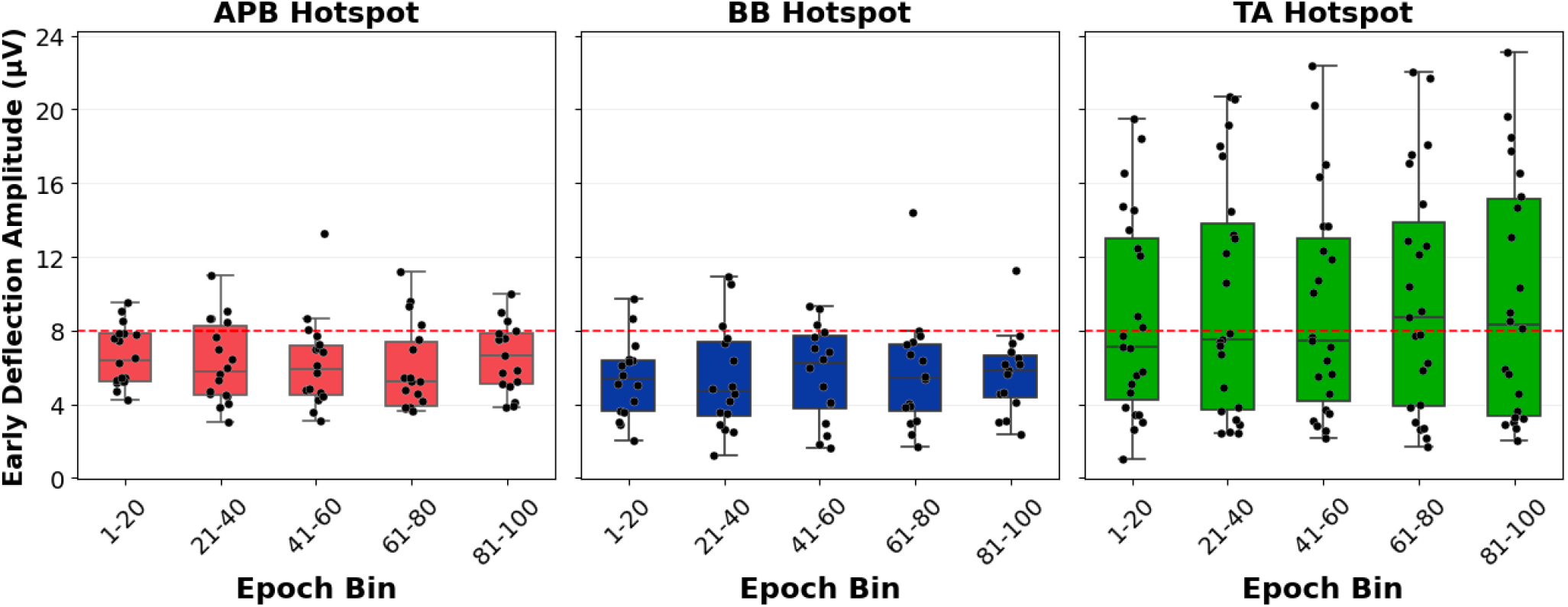
Early TEP stability. 100 TMS-EEG epochs are shown across all subjects. Box: median and interquartile range. Whiskers: furthest data points within 1.5 times interquartile range from the box edges. Dots: individual observations.

There was no statistically significant difference between bins across the TMS-EEG session for any hotspot (Figure 7). Within-subject variability of the early TEP deflection amplitudes was high, especially in the TA hotspot, but the fluctuations did not appear to follow any specific trend (See Figure S5 in Supplementary Information).

In some sessions, the earliest TEP deflections occurred before the “prototypical” N15-P30 (Figure 2). Such earlier deflections were most common at the TA hotspot (14 cases), followed by 10 cases at the BB hotspot, and 7 cases at the APB hotspot. Importantly, the early deflection latencies were consistent across consecutive 20-average bins within sessions.

## 4. Discussion

In this work, we assessed the relationship between TMS cortical and corticospinal excitability across the M1, and the feasibility of online TEP-based cortical excitability assessment. We benchmarked the online early-TEP detectability criterion against homotopic muscle-specific RMTs across hand, arm, and leg M1 representations, and tested the short-term stability of TEP-based cortical excitability. Our main findings were the following:

□ **Both RMT and CET increased from lateral to medial M1, but the CET gradient was shallower.**
**□ The thresholds correlated strongly while retaining hotspot-specificity.**
**□ Online early-TEP thresholding was feasible across M1, but sensitive to electrode selection, deflection latency, and within-day amplitude drift.**

Taken together, these findings indicate that EEG- and EMG-defined thresholds are neither independent nor interchangeable in M1. We believe their joint assessment could form the basis of a **multidimensional motor system excitability profile** integrating local cortical recruitment, muscle-specific descending corticospinal output, and, prospectively, network-level propagation. Below, we discuss our findings and possible future steps toward an integrated dose map, linking online EMG and EEG output signals/readouts along the M1.

### Parallel but shallower latero-medial gradient for CETs compared to RMTs

Our finding extends the established EMG-derived RMT gradient along the M1 (Chen, Yaseen, *et al*., 1998; Yuasa *et al*., 2022; Koponen *et al*., 2024) by showing an analogous gradient for an EEG-derived early TEP deflection threshold (CET). The commonality of these two gradients, represented as both MSO% and induced EF (V/m), suggests that excitability of the stimulated cortical area is the key factor that anatomical depth does not level out. However, the substantially shallower CET gradient suggests that differences in local cortical excitability cannot account for the RMT gradient alone. We suggest that the observed differences in RMTs along M1 reflect differences in cortical excitability, which are amplified by differences in the strength of corticomotoneuronal projections. This is consistent with the view that RMTs reflect the excitability of the corticospinal motor system rather than being a measure of purely cortical origin (Siebner *et al*., 2022). We propose that the shallower intensity-output curves of BB and TA muscles compared to intrinsic hand muscles (Koponen *et al*., 2024) reflect wiring differences in the corticospinal networks. We further suggest that the divergence between the EMG and EEG gradients may reflect relative contributions of muscle-specific corticospinal outputs at the individual level, and might provide a neurophysiological proxy for the ability to individuate muscle forces, but this interpretation should be confirmed in further studies. In particular, it will be important to assess these effects during different motor tasks, such as execution, imagination, observation, which are known to change corticospinal excitability (Chen, Yaseen, *et al*., 1998; Roosink and Zijdewind, 2010; Feurra *et al*., 2019; Bonnesen *et al*., 2022; Grilc *et al*., 2024; Morozova *et al*., 2024; Nieto-Doval *et al*., 2025) and in clinical conditions such as stroke, spinal cord injury, or dystonia.

### Correspondence and hotspot-specificity of the EEG- and EMG-derived thresholds

The strong associations between CETs and RMTs suggest that they share motor system excitability at the individual level, consistent with some of the previous evidence (Mäki and Ilmoniemi, 2010; Gedankien *et al*., 2017). However, the within-subject range in CETs was substantial, arguing against the common use of the hand RMT (Ridding and Rothwell, 1997; Boroojerdi *et al*., 2001; Möller *et al*., 2009) to stimulate targets outside the M1 hand area. Moreover, the EEG- and EMG-derived thresholds showed progressively less concordance toward more medial portions of M1.

A key feature of our study was the use of the compound EMG-EEG hotspots. We did not compromise MEP quality, but, instead, we optimized the targeting to achieve both robust MEPs and artifact-free TEPs for thresholding. Consequently, the resulting threshold correspondence reflects the relationship between cortical and corticospinal excitability under matched stimulation conditions, rather than independently tuned thresholds.

Stimulation orientation strongly influences both MEPs (Brasil-Neto *et al*., 1992; Raffin *et al*., 2015; Reijonen *et al*., 2020; Nazarova, Novikov, *et al*., 2021) and TEPs (Casarotto *et al*., 2010). Thus, future studies should compare the orientation dependence of both thresholds simultaneously. It would be especially interesting to test paired-pulse TMS combined with EEG and EMG, to assess whether CET corresponds to the minimum conditioning-stimulus intensity required to modulate the test response across different paired-pulse TMS phenomena (Fedele *et al*., 2016; Iscan *et al*., 2016; Cash and Ziemann, 2021). Indeed, it has already been reported that conditioning-stimulus intensity can affect TEP and MEP modulation differently (Rawji *et al*., 2021) and that the effects of conditioning pulse orientation vary across paired-pulse phenomena (Tugin *et al*., 2021; Souza *et al*., 2025). So, a combined TMS-EMG-EEG thresholding approach could help clarify the relationship between EEG-detectable cortical activity and the recruitment of circuits that influence corticospinal output.

### Sub-RMT early TEP threshold

We used here an 8-µV criterion in early TEP deflections as an operational CET, instead of inferring cortical excitability from an indirect measure of corticospinal excitability. In most subjects, this criterion was reached below the corresponding RMT, indicating that prominent early TEPs do not require reliable recruitment of the target muscle. This choice of early TEP amplitude criterion agrees with previous studies that adopted early EEG response criteria of 6–8 µV rather than 10 µV (Casarotto *et al*., 2022; Ukharova *et al*., 2025). Our results agree with earlier studies showing that already 60% of RMT and 30-40 V/m were enough to evoke TEPs (Komssi *et al*., 2007; Saari *et al*., 2018). However, we observed no universal relationship between RMTs and CET at the same hotspot, and the rule that RMTs are higher than CET was not absolute for any hotspot we investigated. Accordingly, it may be important to use both EMG and EEG readouts for TMS dosage estimation, depending on whether one is primarily assessing cortical or corticospinal drive.

### Feasibility of online EEG-derived TMS thresholding

Our findings confirm the feasibility of online early-TEP thresholding but also identify several challenges for its standardization. In most cases there were two electrodes with comparable big early TEP deflection, raising the question of how the response selection may be standardized without assuming a fixed scalp topography. Spatial filtering may be an alternative to single-electrode deflection detection (Couto *et al*., 2026), but its suitability for online threshold estimation is unclear.

Second, the latency of the earliest identifiable TEP deflection varied across participants and stimulation sites, with earlier deflections observed more frequently at the TA hotspot. Earlier deflections were not confined to a single site: in more than half of cases, they occurred at two or all three hotspots within the same participant. These findings suggest that both participant- and site-specific factors may contribute to early TEP timing, consistent with reported regional differences in TMS-evoked response dynamics (Passera *et al*., 2022) and the sensitivity of early TEPs to stimulation parameters (Lucarelli *et al*., 2025). Further investigation is needed to distinguish the contributions of participant characteristics, stimulation site, and coil orientation.

Third, obtaining the target amplitude during CET determination did not guarantee that the same intensity would reproduce that amplitude later. We expected that selecting the largest early deflection close to the stimulation site, may preferentially select an upward noise fluctuation (Clayson, Baldwin and Larson, 2013). However, while at the hand representation subsequent early-TEP amplitudes indeed tended to be lower than the calibration target of 8 µV, at the TA representation subsequent amplitudes showed greater between-subject variation, with deviations both above and below the target. One hypothesis is that matching a single amplitude endpoint does not match the sensitivity of the underlying response: input–output curves may differ across M1 (Raffin *et al*., 2020). From a practical standpoint, we need to agree on repeatability limits, assessing absolute agreement and smallest detectable change, as it was partly done for TMS-EMG mapping parameters (Nazarova *et al*., 2023). When the objective is to maintain a consistent evoked response, adaptive intensity re-estimation, real-time TEP visualization, and EEG-guided optimization approaches (Casarotto *et al*., 2022; Tervo *et al*., 2022) could be used to detect deviations that exceed measurement uncertainty.

## 5. Conclusion and Future Directions

The findings of the present study argue against using a hand-muscle RMT to set cortical excitability dosage along M1. Within subjects, CETs and their corresponding muscle-specific RMT differed at all three M1 representations, and this gap widened from arm to leg representations. Online EEG guided dosing in the motor cortex may achieve sufficient early cortical engagement while reducing, although not eliminating, motor reafference and peripheral co-stimulation. EEG guided dosing for the motor cortex can be particularly relevant in clinical populations like stroke, when substantial fraction of patients do not have MEPs at least in some hand muscles (Turton *et al*., 1996; Stinear CM *et al*., 2012; Nazarova, Kulikova, *et al*., 2021); and where contralesional RMT is sometimes used to set ipsilesional stimulation intensity for TMS-EEG (Tscherpel *et al*., 2020).

While we provide a site-by-site comparison of cortical and corticospinal thresholds along M1, future work should determine how movement reweights its components and whether these state-dependent changes explain individual motor function, as movement may reweight cortical and corticospinal readouts (Nikulin *et al*., 2003; Desideri *et al*., 2019). We hypothesize that movement-related changes in cortical excitability will be stimulation-site specific and that the spatial pattern of this reconfiguration will relate to individual behavioral abilities, such as hand dexterity and its problems and recovery in clinical conditions like stroke or dystonia.

## Acknowledgements

Authors are deeply grateful to Zhenrui Cheng for her early support in data collection and analysis, and Matilda Montonen for her support at the initial stages of data collection. Authors are very grateful to all our participants. This project has received funding from the European Union’s Horizon Europe research and innovation programme under the Marie Skłodowska-Curie grant agreement No 101060584, and the Research Council of Finland.

## Data availability statement

Data supporting the findings of this study are available from the corresponding author upon reasonable request.

