## Supplementary Figures and Tables for "Matching Corticospinal and Cortical Excitability along the motor cortex: TMS-EMG-EEG study"

### Supplementary Information

### Methods

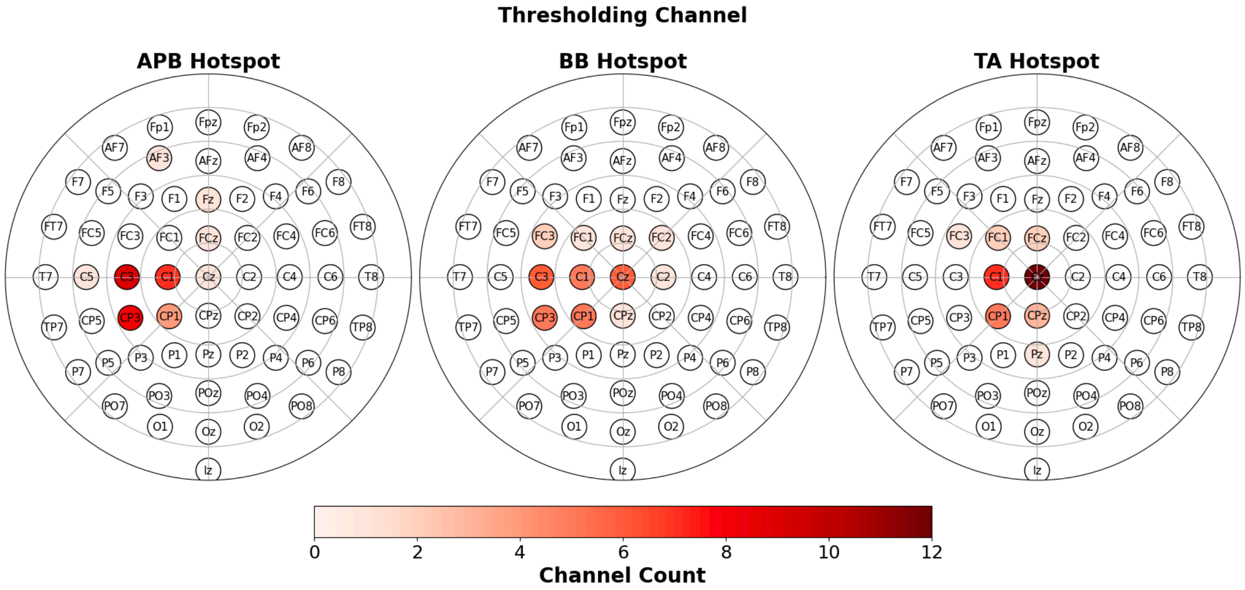

**Figure S1.** *Channels used for CET determination for all three hotspots.*

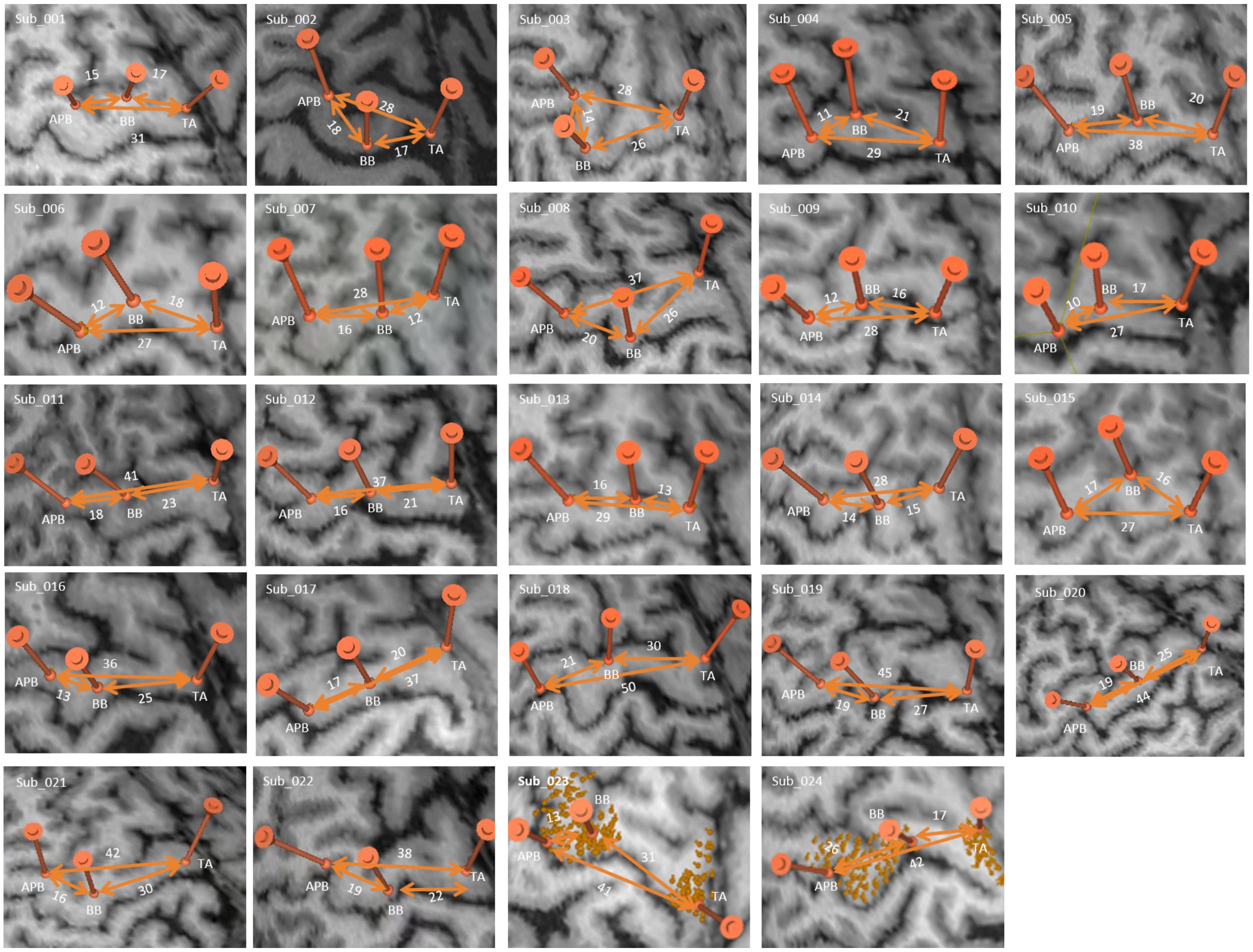

***Figure S2:*** *Locations of all three compound hotspots on the primary motor cortex (the three orange circles) for APB (left), BB (middle), and TA (right) in the 24 analyzed subjects, with Euclidean distances in mm. *The shortest distance between the APB and BB spots was still greater than 10 mm. The last two subjects include examples of the rough mapping with the hotspots.*

### Results

| | Threshold Descriptive Statistics V/m | | | | | |
|---|---|---|---|---|---|---|
| | RMT | | | CET | | |
| | APB | BB | TA | APB | BB | TA |
| Number of values | 24 | 24 | 24 | 24 | 23 | 24 |
| Minimum | 42.5 | 54.5 | 63.5 | 34 | 39.5 | 46 |
| Median | 61 | 76 | 95.5 | 54.5 | 61.5 | 72.75 |
| Maximum | 158.5 | 165 | 155 | 104 | 116 | 127 |
| Range | 116 | 110.5 | 91.5 | 70 | 76.5 | 81 |
| Mean | 67.66 | 78.25 | 100.6 | 60.71 | 66.81 | 76.83 |
| Std. Deviation | 24.38 | 24.15 | 25.9 | 17.14 | 16.35 | 18 |
| Std. Error of Mean | 4.977 | 4.93 | 5.287 | 3.498 | 3.41 | 3.675 |
| Coefficient of variation | 36.04% | 30.86% | 25.74% | 28.22% | 24.47% | 23.43% |

***Table S1:*** *Descriptive Statistics of RMT and CET thresholds (V/m)*

| | Threshold Descriptive Statistics %MSO | | | | | |
|---|---|---|---|---|---|---|
| | RMT | | | CET | | |
| | APB | BB | TA | APB | BB | TA |
| Number of values | 24 | 24 | 24 | 24 | 23 | 24 |
| Minimum | 32 | 28 | 40 | 30 | 49 | 41 |
| Median | 44 | 54 | 70 | 41 | 48 | 57 |
| Maximum | 83 | 64 | 87 | 65 | 100 | 81 |
| Range | 51 | 36 | 47 | 35 | 51 | 40 |
| Mean | 47.42 | 43.08 | 56.25 | 48.87 | 73.25 | 58.58 |
| Std. Deviation | 11.07 | 7.94 | 11.87 | 8.498 | 14.24 | 9.339 |
| Std. Error of Mean | 2.26 | 1.621 | 2.424 | 1.772 | 2.907 | 1.906 |
| Coefficient of variation | 23.35% | 18.43% | 21.11% | 17.39% | 19.44% | 15.94% |

***Table S2:*** *Descriptive Statistics of RMT and CET thresholds (%MSO)*

| Thresholds in V/m | | | | | | | | | |
|---|---|---|---|---|---|---|---|---|---|
| | APB | | | BB | | | TA | | |
| Subject | rMT | CET | % Diff | rMT | CET | % Diff | rMT | CET | % Diff |
| Sub_001 | 55 | 50 | 5 | 60.5 | 61.5 | -1 | 63.5 | 64 | -0.5 |
| Sub_002 | 53.5 | 54.5 | -1 | 54.5 | 58.5 | -4 | 78 | 72.5 | 5.5 |
| Sub_003 | 98 | 88 | 10 | 115 | 81.5 | 33.5 | 129 | 108.5 | 20.5 |
| Sub_004 | 47.5 | 50 | -2.5 | 55.5 | 57.5 | -2 | 76 | 60 | 16 |
| Sub_005 | 84.5 | 60 | 24.5 | 85 | 61.5 | 23.5 | 106.5 | 66 | 40.5 |
| Sub_006 | 54.4 | 54.5 | -0.1 | 77 | X | | 88 | 59 | 29 |
| Sub_007 | 61 | 57.5 | 3.5 | 78 | 59 | 19 | 80.5 | 70 | 10.5 |
| Sub_008 | 72.5 | 56.5 | 16 | 75 | 61.5 | 13.5 | 65.5 | 69 | -3.5 |
| Sub_009 | 53.5 | 53.5 | 0 | 66 | 65.5 | 0.5 | 84 | 73 | 11 |
| Sub_010 | 64 | 56 | 8 | 58 | 65 | -7 | 111 | 60 | 51 |
| Sub_011 | 84.5 | 79 | 5.5 | 72 | 71 | 1 | 88 | 75 | 13 |
| Sub_012 | 64 | 53.5 | 10.5 | 80 | 53.5 | 26.5 | 103 | 82 | 21 |
| Sub_013 | 48.5 | 50 | -1.5 | 80 | 56 | 24 | 97 | 64 | 33 |
| Sub_014 | 67.5 | 68 | -0.5 | 88 | 83 | 5 | 126 | 87.5 | 38.5 |
| Sub_015 | 52.5 | 53 | -0.5 | 60 | 60 | 0 | 95.5 | 88.5 | 7 |
| Sub_016 | 158.5 | 104 | 54.5 | 165 | 116 | 49 | 155 | 127 | 28 |
| Sub_017 | 68.5 | 80 | -11.5 | 93 | 83 | 10 | 116 | 84.5 | 31.5 |
| Sub_018 | 51.5 | 34 | 17.5 | 57.5 | 39.5 | 18 | 95.5 | 72 | 23.5 |
| Sub_019 | 42.5 | 42 | 0.5 | 61 | 51 | 10 | 72 | 46 | 26 |
| Sub_020 | 94 | 98 | -4 | 97 | 94 | 3 | 150.5 | 93.6 | 56.9 |
| Sub_021 | 50.5 | 53 | -2.5 | 55 | 60 | -5 | 90 | 82.6 | 7.4 |
| Sub_022 | 77.5 | 61.6 | 15.9 | 89 | 75.7 | 13.3 | 117 | 91.7 | 25.3 |
| Sub_023 | 61 | 52 | 9 | 85 | 69.5 | 15.5 | 144 | 91 | 53 |
| Sub_024 | 59 | 48.5 | 10.5 | 71 | 53 | 18 | 84 | 56.5 | 27.5 |
| Total CE-RMT > RMT: | | | 9 | | | 5 | | | 2 |

***Table S3:*** *Individual subject threshold values for RMT and CET in V/m.*

| Thresholds in %MSO | | | | | | | | | |
|---|---|---|---|---|---|---|---|---|---|
| | APB | | | BB | | | TA | | |
| Subject | rMT | CET | % Diff | rMT | CET | % Diff | rMT | CET | % Diff |
| Sub_001 | 44% | 40% | 4% | 47% | 49% | -2% | 59% | 60% | -1% |
| Sub_002 | 47% | 48% | -1% | 51% | 55% | -4% | 72% | 67% | 5% |
| Sub_003 | 53% | 50% | 3% | 63% | 46% | 17% | 79% | 65% | 14% |
| Sub_004 | 38% | 40% | -2% | 45% | 47% | -2% | 69% | 56% | 13% |
| Sub_005 | 53% | 38% | 15% | 61% | 45% | 16% | 84% | 53% | 31% |
| Sub_006 | 39% | 39% | 0% | 50% | X | X | 63% | 49% | 14% |
| Sub_007 | 43% | 40% | 3% | 63% | 48% | 15% | 66% | 62% | 4% |
| Sub_008 | 51% | 39% | 12% | 59% | 49% | 10% | 61% | 64% | -3% |
| Sub_009 | 44% | 44% | 0% | 53% | 51% | 2% | 63% | 55% | 8% |
| Sub_010 | 41% | 38% | 3% | 42% | 42% | 0% | 74% | 45% | 29% |
| Sub_011 | 52% | 48% | 4% | 52% | 51% | 1% | 64% | 55% | 9% |
| Sub_012 | 49% | 41% | 8% | 59% | 43% | 16% | 82% | 60% | 22% |
| Sub_013 | 44% | 42% | 2% | 74% | 51% | 23% | 91% | 58% | 33% |
| Sub_014 | 50% | 52% | -2% | 69% | 61% | 8% | 100% | 70% | 30% |
| Sub_015 | 32% | 34% | -2% | 40% | 40% | 0% | 59% | 55% | 3% |
| Sub_016 | 83% | 56% | 27% | 87% | 63% | 24% | 91% | 72% | 19% |
| Sub_017 | 38% | 45% | -7% | 55% | 51% | 4% | 67% | 55% | 12% |
| Sub_018 | 43% | 28% | 15% | 45% | 30% | 15% | 71% | 52% | 19% |
| Sub_019 | 36% | 34% | 2% | 52% | 42% | 10% | 58% | 41% | 17% |
| Sub_020 | 59% | 64% | -5% | 66% | 65% | 1% | 96% | 63% | 33% |
| Sub_021 | 40% | 44% | -4% | 45% | 47% | -2% | 75% | 69% | 6% |
| Sub_022 | 66% | 53% | 13% | 74% | 64% | 10% | 100% | 81% | 19% |
| Sub_023 | 56% | 41% | 15% | 57% | 44% | 13% | 65% | 53% | 12% |
| Sub_024 | 37% | 36% | 1% | 41% | 40% | 1% | 49% | 46% | 3% |
| Total CE-RMT > RMT: | | | 7 | | | 4 | | | 2 |

***Table S4:*** *Individual subject threshold values for RMT and CET in %MSO.*

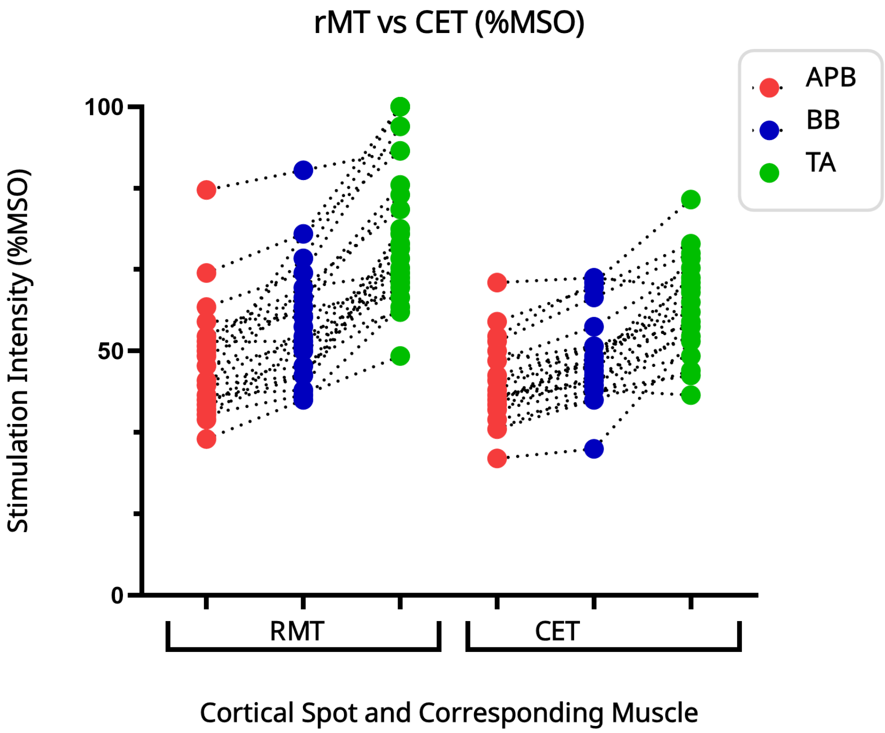

***Figure S3:*** *RMTs vs. CETs (%MSO) gradient at the hand (APB; red), arm (BB; blue), and leg (TA; green) hotspots on the primary motor cortex.*

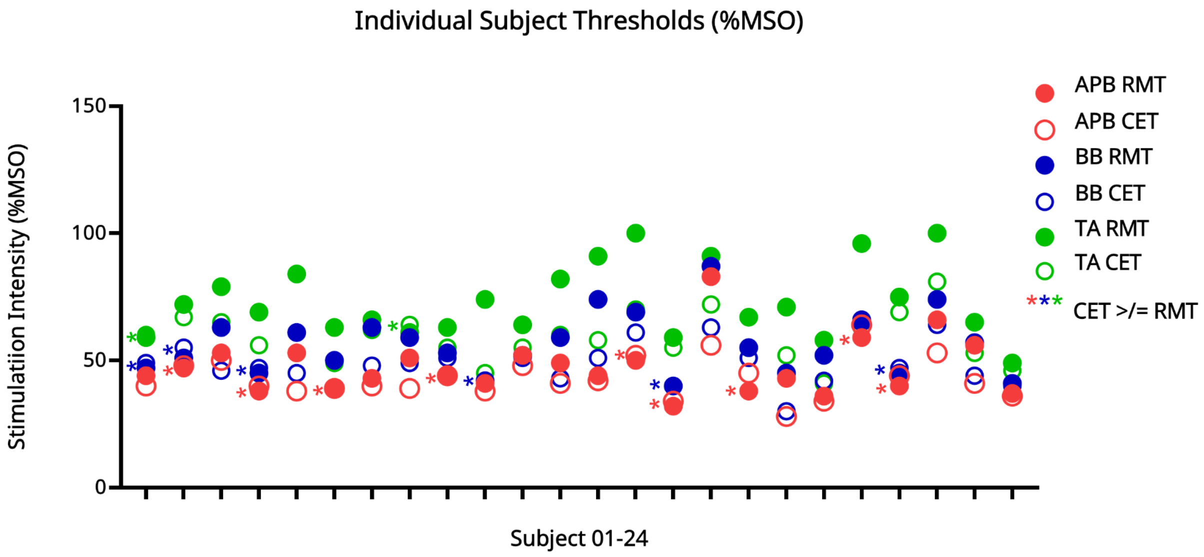

***Figure S4:*** *Subject-specific RMTs vs. CETs (%MSO) at hand (APB, red), arm (BB, blue) and leg (TA, green) hotspots on the primary motor cortex. Filled circles represent RMTs, empty circles represent CETs. * indicates cases where the CET was greater than or equal to RMT at the same spot (according to the spot color coding).*

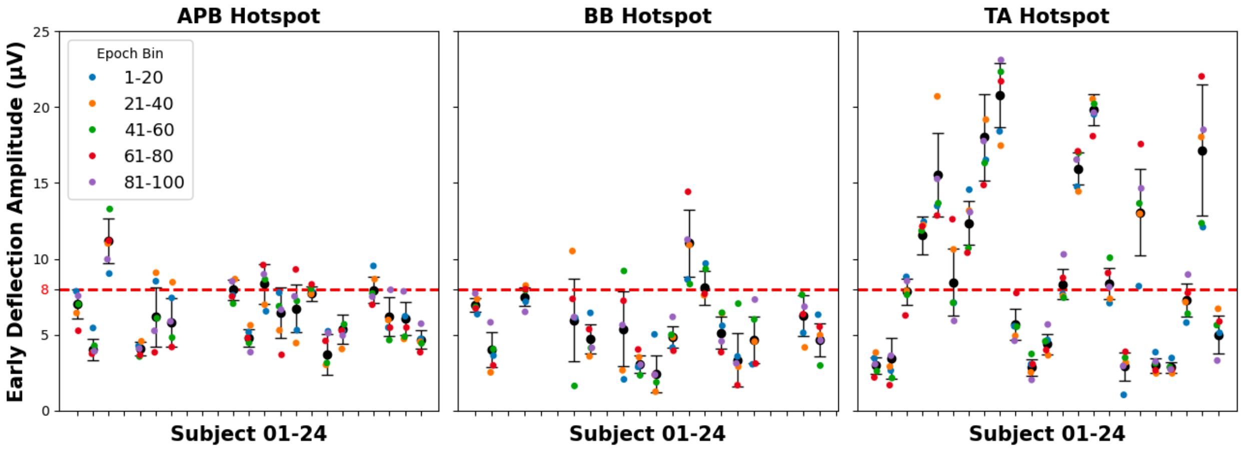

***Figure S5:*** *N15-P30 peak-to-peak values (mean of 20 trials) per subject during rest for each of the muscles addressed.*
